# Using high-throughput phenotypes to enable genomic selection by inferring genotypes

**DOI:** 10.1101/2020.02.28.969600

**Authors:** Andrew Whalen, Chris Gaynor, John M Hickey

**Affiliations:** The Roslin Institute and Royal (Dick) School of Veterinary Studies, The University of Edinburgh, Midlothian, Scotland, UK

**Author notes:** Corresponding author: AW. Email addresses: CG, JMH.

## Abstract

In this paper we develop and test a method which uses high-throughput phenotypes to infer the genotypes of an individual. The inferred genotypes can then be used to perform genomic selection. Previous methods which used high-throughput phenotype data to increase the accuracy of selection assumed that the high-throughput phenotypes correlate with selection targets. When this is not the case, we show that the high-throughput phenotypes can be used to determine which haplotypes an individual inherited from their parents, and thereby infer the individual’s genotypes. We tested this method in two simulations. In the first simulation, we explored, how the accuracy of the inferred genotypes depended on the high-throughput phenotypes used and the genome of the species analysed. In the second simulation we explored whether using this method could increase genetic gain a plant breeding program by enabling genomic selection on non-genotyped individuals. In the first simulation, we found that genotype accuracy was higher if more high-throughput phenotypes were used and if those phenotypes had higher heritability. We also found that genotype accuracy decreased with an increasing size of the species genome. In the second simulation, we found that the inferred genotypes could be used to enable genomic selection on non-genotyped individuals and increase genetic gain compared to random selection, or in some scenarios phenotypic selection. This method presents a novel way for using high-throughput phenotype data in breeding programs. As the quality of high-throughput phenotypes increases and the cost decreases, this method may enable the use of genomic selection on large numbers of non-genotyped individuals.

## Introduction

In this paper we develop and test a method which uses high-throughput phenotypes to infer the genotypes of an individual. The inferred genotypes can then be used to perform genomic selection. The routine use of genomic selection in plant breeding programs has increased the rate genetic gain in many crop species by increasing the accuracy of selection. In order to perform genomic selection, genotypes must be available on the individuals evaluated. In many breeding programs it is still prohibitively expensive to genotype all of the selection candidates, particularly when a large number of individuals need to be evaluated. In these situations, selection is done either at random, or by using low-accuracy proxies, such as visual selection [1]. High-throughput phenotypes offer one way to increase the accuracy of selection on non-genotyped individuals by providing accurate proxies for selection targets [2–4]. High-throughput phenotypes, often based on spectral data such as near infrared spectrometry (NIR), can be collected cheaply and non-destructively on a large number of individuals [5–7]. Similar data types are also potentially available in animal breeding, for example from the routine collection of milk infrared spectrometry data in dairy cattle [8].

Using high-throughput phenotypes to increase the accuracy of selection has primarily focused on identifying phenotypes that correlate with the selection targets and using them either as proxies for selection targets for non-genotyped individuals [5,6,9], or as correlated traits in genomic prediction models for genotyped individuals [3,10]. In some cases, high-throughput phenotypes may not correlate with selection targets. This will be particularly the case when the individual’s growing environment is dissimilar from selection environment, such as when individuals are grown in a greenhouse or winter nursery. In these cases, it is important to develop methods that allow high-throughput phenotypes to be used to increase the accuracy of selection for non-genotyped individuals.

One approach is to use the high-throughput phenotypes as a stand-in for genomic markers, as in a method that Rincent et al. called phenomic selection [11]. Phenomic selection exploits the fact that with a large number of heritable high-throughput phenotypes, the phenotypic covariance between individuals is close to the genetic covariance between individuals. This changes the prediction problem from one of detecting relationships between high-throughput phenotypes and selection targets, to one of estimating the relationships between individuals. Rincent et al. demonstrated that phenomic selection could be used to predict selection targets in both the environment where the high-throughput phenotypes were collected, and in a separate environment where the plants were grown under different management conditions.

Phenomic selection is an attractive way to increase the accuracy of selection on non-genotyped individuals. However, it does not exploit existing genotype and phenotype data, and deploying phenomic selection in practice would require developing an additional training population with both selection targets and the same set of high-throughput phenotypes measured.

An alternative way to use high-throughput phenotypes to increase the accuracy of selection is to use the high-throughput phenotypes to infer the genotypes of non-genotyped individuals, and thus enable genomic selection on these individuals. This approach has two advantages. First, the inferred genotypes can be integrated into existing genomic selection frameworks, taking advantage of existing genomic training populations. Second, once the genotypes are inferred, the genomic predictions produced will be independent of the individual’s growing environment, or the particular set of high-throughput phenotypes measured.

Here, we present a method for using high-throughput phenotypes to infer the genotypes of non-genotyped individuals. To do this, we extend research on genetic imputation which uses long shared haplotype segments to impute genotypes from sparse SNP array, sequence, or genotyping-by-sequencing data [12–14]. Imputation can be greatly simplified by using family and pedigree data to reduce the pool of haplotypes under consideration [15–17]. In a bi-parental cross, only the parental haplotypes need to be considered which results in high-accuracy imputation even when few markers are used [16,18,19].

Instead of using sparse genetic data, it may be possible to use phenotypic information to infer which genotypes an individual carries, by evaluating which combinations of inherited haplotypes are likely to give rise to the observed phenotypes. We implement this technique by representing haplotype combinations as a series of segregation states, which give the parent of origin for the haplotype at each locus. We then sequentially sample the segregation states for each chromosome. The sampling process is guided by comparing the expected genetic value for the individual, conditional on the sampled segregation state, to the observed high-throughput phenotypes.

In this paper, we first describe how we use high-throughput phenotypes to infer genotypes. We then use simulations to explore two questions. First, how the accuracy of the inferred genotypes depended on the high-throughput phenotypes used and the genome of the species analysed. Second, whether using this method could increase genetic gain in a plant breeding program by enabling genomic selection on non-genotyped individuals. In the first simulation we found that genotype accuracy was higher if more high-throughput phenotypes were used and if those phenotypes had higher heritability. We also found that genotype accuracy decreased as the genome of the species increased. In the second set of simulations we found that the inferred genotypes could be used to enable genomic selection on non-genotyped individuals and increase genetic gain compared to random selection, or in some scenarios phenotypic selection. This method could have value particularly in cases where selection targets are hard to measure in a plant’s growing environment.

## Materials and Methods

### Using high-throughput phenotypes to infer genotypes

In this method, we use high-throughput phenotypes to determine which haplotypes an individual inherited from their parents, and thereby infer the individual’s genotypes. An individual in this context refers to either an inbred plant or a group of genetically identical plants taken from the inbred line. To perform inference, we iteratively sample the segregation state at each locus on each chromosome based on how well the expected genetic value for a segregation state matches the observed high-throughput phenotypes. The segregation state indicates whether the individual inherits the paternal or maternal haplotype at that locus [20].

This method takes as inputs: (i) single nucleotide polymorphism (SNP) array genotypes of an individual’s parents, (ii) high-throughput phenotypes on the individual, and (iii) SNP marker effects for each genotyped locus and each high-throughput phenotype. The method outputs inferred genotype dosages.

We assume the parents and offspring are part of a bi-parental cross, and both the parents and offspring are fully inbred. This method could be adapted to animals or outbred plants if the genotypes of the parents are known and phased.

This method can be broken down into three pieces. First, how segregation states can be translated into expected genetic values. Second, how to sample the segregation states for a single chromosome. Third, how to sample the entire genome, and how to translate the sampled segregation states into inferred genotype dosages.

### Converting segregation states into genetic values

We convert segregation states into genetic values by first using the genotypes of the parents to convert the segregation states to offspring genotypes, and then using the SNP effect estimates to convert the genotypes into a genetic value for each high-throughput phenotype.

We assume an additive model for each high-throughput phenotype, *j*:

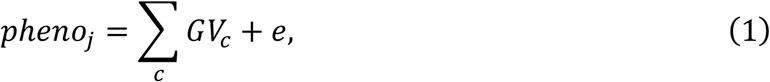

where the observed phenotype is modelled as a summation of the genetic values for each chromosome, *c*, combined with an environmental random effect term, e∼*N*(0, *σ*_*e*_). The genetic value for a particular chromosome is given by:

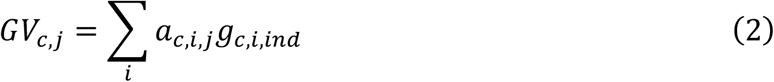

where *a*_*c,i,j*_, gives the SNP effect for locus *i*, and phenotype *j*, and *g*_*c,i,ind*_ gives the genotype for individual *ind*.

For an individual with unknown genotypes, we use the genotypes of their parents to translate segregation states into genetic values. We define the segregation state, *x*_*c,i*_ ∈ {0,1}, as the parent of origin for the individual’s genotype at locus, *i*, on chromosome, *c.* Given a series of segregation states, the resulting genetic value is:

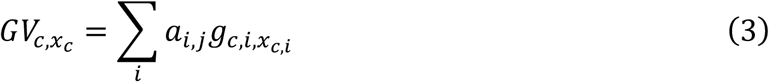

where 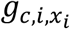 is the genotype of the father if *x*_*c,i*_ is 0, or the genotype of the mother if *x*_*c,i*_is 1.

### Sampling segregation states for a single chromosome

We sample the segregation states on a particular chromosome by first estimating a residual phenotype, and then using the residual phenotype to guide the sampling of segregation states. The residual phenotype for a particular chromosome is defined as observed phenotype minus the genetic value for the remaining chromosomes:

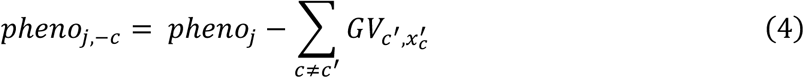

Segregation states are then sampled sequentially from the first locus to the last locus. At each locus, the segregation state, *x*_*c,i*_ ∈ {0,1}, is sampled proportional to:

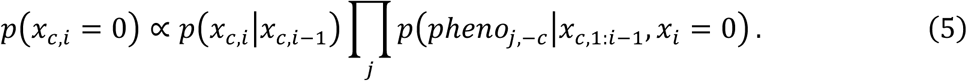

The first term, *p*(*x*_*c,i*_|*x*_*c,i*−1_), is the probability of a segregation state conditional on the previous segregation state. The second term, *p*(*pheno*_*j*,−*c*_|*x*_*c*,1:*i*−1_, *x*_*i*_ = 0), is the probability residual phenotype conditional on a segregation state.

For the first term we assume that the sequence of segregation states follows a Markov process with a recombination rate, *r*. This gives:

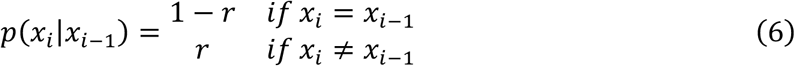

For second term we use a normal approximation to approximate the expected genetic value for the unsampled portion of the chromosome (from loci *i* up to *n*):

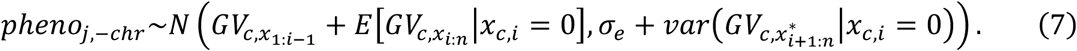

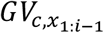 is the genetic value for the previously sampled loci on the chromosome. 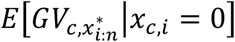 is the expected genetic value for the remaining loci on the chromosome, conditional on *x*_*c,i*_ = 0. 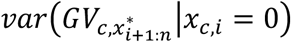 is the variance of the genetic value for the remaining loci on the chromosome conditional on *x*_*c,i*_ = 0. *σ*_*e*_ is the environmental variance.

We recursively calculate expected genetic value using the relationship:

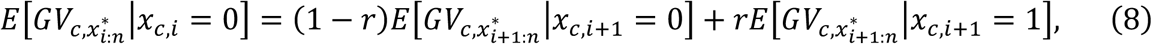

and recursively calculate the variance using the relationship:

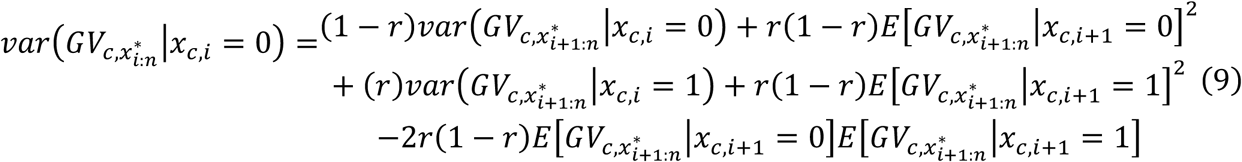

### Sampling segregation states for multiple chromosome and inferring genotype dosages

We performed sampling for the entire genome in a series of independent Markov chains. In each chain, we initialized the genetic values for each chromosomes as their mean genetic value (assuming each segregation state was equally likely). We then sequentially sampled each chromosome in the genome using the single chromosome sampling method described above. We repeated this sampling process for 100 steps, in each step the segregation state for each chromosome was re-sampled once. This entire process was repeated to create 10 independent chains. Results were averaged across all steps and chains.

The use of multiple steps and chains allows for a broader exploration of likely segregation states. Following previous research into Markov Chain Monte Carlo sampling, which found that averaging across samples produced more accurate results than first thinning the samples to reduce autocorrelation, we did not thin the samples [21]. Preliminary simulations found that the accuracy did not increase across steps, so a burn-in period was not used.

We calculated the inferred genotype dosage as the mean genotype averaged across all steps and chains

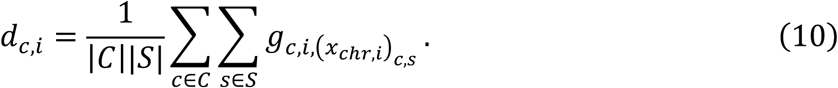

The summation is over each chain, *c* ∈ *C*, and each step, *s* ∈ *S*, and 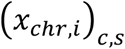 is the segregation state sampled in step *s*, and chain *c*.

### Simulations

We tested this method in two sets of simulations. First, we performed a parameter sensitivity analysis to quantify how the accuracy of the inferred genotypes depended on the high-throughput phenotypes used and the genome of the species analysed. Second, we simulated a plant breeding program to see whether using this method could increase genetic gain by enabling genomic selection on non-genotyped individuals. Both simulations shared a common genetic architecture, phenotypic architecture, and method for inferring SNP effects, but differed in the population structure and scenarios analysed.

### Genetic Architecture

In both simulations, we simulated a genome consisting of between 5-21 chromosomes for a hypothetical plant species. Founder genomes were generated using the Markovian coalescent simulator, MaCS [22], using a population history similar to wheat. In MaCS each chromosome had a physical length of 8×10^8^ base pairs, with a recombination rate depending on genome size (between 5×10^−9^ and 2×10^−8^ recombinations per base pair), and a mutation rate of 2×10^−9^ per base pairs. Effective population size was set to 50, with linear piecewise increases to 1,000 at 100 generations ago, 6,000 at 1,000 generations ago, 12,000 at 10,000 generations ago, and 32,000 at 100,000 generations ago. The values were chosen to follow the evolution of effective population size in wheat [23].

The founder haplotypes were dropped through the population using AlphaSimR [24]. In AlphaSimR, we extracted between 3,100 and 4,000 segregating sites per chromosome. Of the segregating sites, 3,000 sites were used as potential quantative trait loci (QTL), and the remaining 100 to 1,000 sites were used form a SNP array. The segregating sites used to create the SNP array did not overlap with the segregating sites used as QTL. We did not simulate any genotyping errors, and assumed there were no residual heterozygous loci for doubled haploid individuals.

### Phenotypic architecture

Phenotypes were simulated using an additive genetic model. For each phenotype, 100 QTL per chromosome were randomly chosen from the pool of 3,000 potential QTL per chromosome. QTL effects were sampled from a standard normal distribution, N(0,1), and linearly scaled to produce genetic values with a target genetic variance in the founder generation. Each individual’s genetic value was calculated by summing their QTL effects across the genome, and their phenotypes calculated as the sum of their genetic value and a random environmental effect, sampled from a standard normal distribution.

In both the parameter sensitivity analysis simulations, and the breeding program simulations, 100 high-throughput phenotypes were simulated. All of the high-throughput phenotypes had the same initial heritability which depended on the scenario analysed.

In the breeding program simulations, we followed the same process to simulate an additional yield phenotype with an initial genetic variance of 0.1 and an environmental variance of 0.4.

### Estimating SNP effects for high-throughput phenotypes

In order to infer genotypes, we estimated SNP effects for each marker on the SNP array. These effects were estimated using ridge regression [25], and assumed that a training population existed which were evaluated on each of the high-throughput phenotypes and the SNP array genotypes. The details of the training population depended on the simulation

To calculate SNP effects, we assumed that the phenotypes followed an additive model:

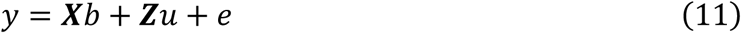

where *y* is a vector of observed phenotypes, ***X*** is a fixed effect design matrix, *b* is a vector of fixed effects, ***Z*** is a design matrix for the random effects, which contains the SNP dosages for each individual at each locus, *u* is a vector of SNP effects, and *e* is a random error term. In both the parameter sensitivity analysis simulations and in the breeding program simulations, we assumed that the training population was grown in the same environment (location and year) as the testing population. Because of this we did not include location as a fixed effect, and the fixed effect design matrix only included the intercept term. This model was fit using the “solveRRBLUP” method in the R package AlphaMME (https://bitbucket.org/hickeyjohnteam/alphamme/).

In the breeding program simulations we also estimated SNP effects and genomic predictions for yield. This estimation process is described in more detail in the breeding program simulations.

### Simulation 1: Parameter Sensitivity Analysis

In Simulation 1, we performed a parameter sensitivity analysis to quantify how the accuracy of the inferred genotypes depended on the high-throughput phenotypes used and the genome of the species analysed. The population design was a series of bi-parental crosses. In the scenarios we varied: (i) the number of high-throughput phenotypes; (ii) the heritability of each phenotype; (iii) the number of markers on the SNP array; (iv) the size of the training population used to estimate the QTL effects; (v) number of chromosomes in the genome; and (vi) the genetic map length for each chromosome.

### Population Design

In this simulation, a target population was created by randomly crossing 100 double haploid parents to produce 100 doubled haploid offspring (100 crosses, 1 individual per cross). A separate training population, was created by randomly crossing the same set of parents to produce 1,000 doubled haploid offspring (1,000 crosses, 1 individual per cross).

We assumed that genotypes were collected on both the parents and the training population, and that high-throughput phenotypes were measured on both the training population and the target population. SNP effects were estimated using either the entire training population, or a subset of the training population (depending on scenario).

### Scenarios

Parameter sensitivity analysis was performed by constructing a base scenario and then varying five parameters one at a time. In the base scenario, we assumed individuals had 10 chromosomes, which were each 100cM in length. We assumed that the SNP array had 500 markers per chromosomes. We assumed that the heritability of each high-throughput phenotype was 0.5, and that training population consisted of all 1,000 individuals. We then varied: (i) the genetic map length between 50cM, 100cM, 200cM, and 400 cM, (ii) the number of chromosomes between 5, 10, 15, and 20, (iii) the number of markers on the SNP array between 100, 500, and 1,000, (iv) the heritability of high-throughput phenotypes between 0.1, 0.2, 0.3, 0.4, 0.5, 0.6, and 0.7, (v) the number of individuals used to train the model between 100, 250, 500, 750, and 1,000 individuals.

In all scenarios, we varied the number of high-throughput phenotypes used. We used either the first 1, 5, 10, 25, 50, or 100 high-throughput phenotypes.

### Assessment

Genotype accuracy was measured by the correlation between an individual’s inferred genotype dosages and their true genotype, corrected by the parent-average genotype [26]. Each scenario was replicated 10 times. Genotype accuracy was averaged across replicates.

### Simulation 2: Breeding program simulations

In Simulation 2, we simulated a plant breeding program to see whether using this method could increase genetic gain by enabling genomic selection on non-genotyped individuals. The breeding program was based on a wheat breeding and the inferred genotypes were used to perform genomic selection on non-genotyped individuals in the headrow. We evaluated a number of scenarios which varied where the headrow was grown, how individuals were selected from the headrow, the species and number of chromosomes simulated (either 7 for barley or 21 for wheat), the number of high-throughput phenotypes used, and the heritability of those phenotypes.

### Breeding program design

The breeding program design was based on the wheat breeding program described in [27]. In this breeding program, a set of inbred lines were used to create a large number of doubled haploid crosses. These crosses were selected in a series of stages, first in an initial headrow, then a preliminary yield trial (PYT), an advanced yield trail (AYT), and two elite yield trails (EYT).

We considered two different structures for the breeding program. In Scenario 1 the headrow was grown in a field in Year 3. In Scenario 2 the headrow was grown in a greenhouse at the end of Year 2, enabling the PYT to be grown in Year 3. Table 1 presents a year-by-year outline of both scenarios, which are described in more detail below.

**Table 1:**
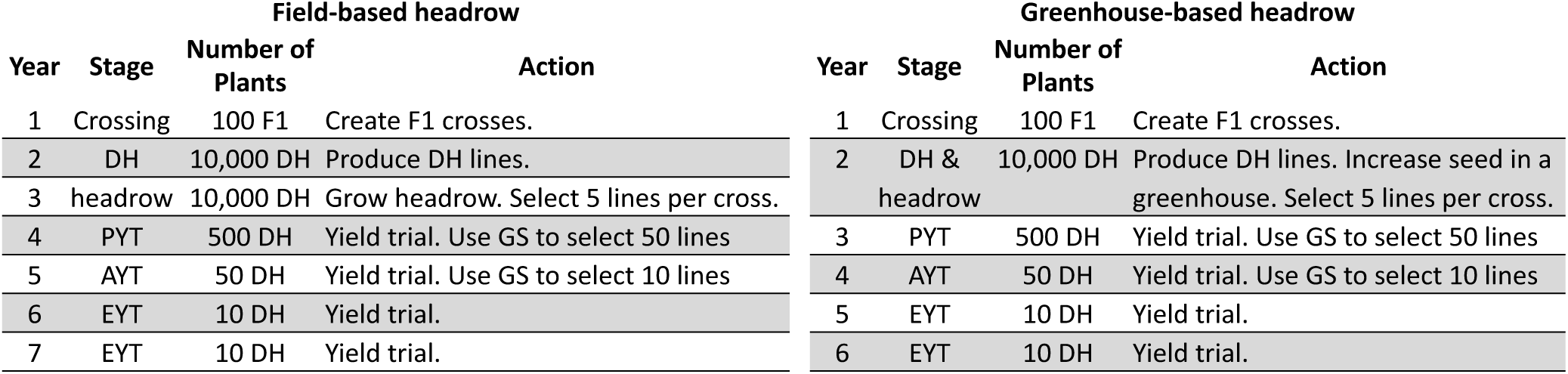
A year-by year description of scenarios 1 and 2. The headrow selection method is used to perform selection in Year 3 (Scenario 1) and Year 2 (Scenario 2). Genomic section (GS) is used to perform selection in future years.

In both scenarios we simulated 20 years of phenotypic selection as a burn-in followed by 20 years of genomic selection.

### Scenario 1: Field-based headrow

#### Year 1

70 parental lines were randomly crossed to produce 100 F_1_ plants. The F_1_ plants were then used to generate 10,000 haploid plants (100 plants per F_1_).

#### Year 2

The genomes of the haploid individuals were doubled to form 10,000 doubled haploid lines.

#### Year 3

Seed from the doubled haploid lines were planted in the field to form a headrow. High-throughput phenotypes were collected on these lines.

500 lines from the headrow (5 per cross) were selected. Headrow selection was done either at random, by visual selection based on the observed yield phenotype, by phenomic selection using the high-throughput phenotypes, by genomic selection using the inferred genotypes, or by genomic selection using the individual’s true genotypes. We describe each selection method in more detail below.

The lines selected from the headrow were genotyped to enable genomic selection in future years.

#### Year 4

The selected lines from the headrow were grown and evaluated in the PYT. We assumed that the PYT was grown in the same field as the headrow from the proceeding year. High-throughput phenotypes were collected on these lines to create a training population to estimate SNP effects for high-throughput phenotype. Genomic selection was used to select 50 lines.

#### Year 5

The 50 selected lines from the PYT were grown and evaluated in the AYT. Genomic selection was used to select 10 lines.

#### Year 6-7

The 10 lines from the AYT were grown and evaluated in the EYT.

#### Year 8

The line with best average performance over Year 6 and Year 7 was released as a variety.

### Scenario 2: Greenhouse-based headrow

The greenhouse-based headrow followed closely to the field-based headrow, except that at the end of Year 2, the doubled haploid plants were grown to increase seed in the greenhouse. These individuals formed a greenhouse-based headrow, and the selected lines were grown in a field-based PYT in Year 3. This scenario evaluates a situation where genomic information is used to reduce the generation interval of a breeding program by removing the need to grow the headrow in the field.

#### Year 1

70 parental lines were randomly crossed to produce 100 F_1_ individuals. The F_1_ plants were then used to generate 10,000 haploid plants (100 plants per F_1_).

#### Year 2

The genomes of the haploid plants were doubled to form 10,000 doubled haploid lines. These lines were then grown in a greenhouse to increase seed forming a greenhouse-based headrow. We assumed that the 500 lines from the PYT were also grown in the greenhouse to act as a training population to estimate SNP effects. High-throughput phenotypes were collected, and a headrow selection method (described in more detail below) was used to select 500 lines (5 per cross). The selected lines were genotyped to enable genomic selection in future years.

#### Year 3-6

The selected lines from Year 2 were grown in a field-based PYT in Year 3. The remainder of Years 3-6 were identical to Years 4-7 in the field-based headrow scenario. In Year 7, the top performing line was released as a variety.

### Parent Selection

In both scenarios, the pool of potential parents each year was created from the combination of the 500 individuals selected from the headrow, the 50 individuals selected from the PYT, the 10 individuals selected from the AYT, and the 10 individuals in both of the EYTs. This created a pool of 570 individuals. Genomic selection was used to select 70 parents to produce crosses for Year 1 of the next generation.

### Burn-in

In both scenarios, we simulated an initial 20 years of burn-in to represent historical breeding. In the burn-in period, individuals in all stages were selected using phenotypic selection. The parents were for each generation were the 50 lines in the AYT and the 10 lines in each EYT (70 total).

### Genomic selection

Genomic selection for yield was performed by using ridge-regression to estimate SNP effects, and then using the SNP effects to calculate estimated genetic values for each line under selection.

To estimate SNP effects, we assumed that the training population consisted of the lines in the PYT, AYT, and EYT from the past 5 years. We assumed these lines were genotyped on a SNP array, and had yield phenotypes collected. For each yield trial, we varied the number of effective phenotypic replicates to represent increased accuracy from multilocation yield trials. The number of effective replicates was 1 in the PYT, 4 in the AYT, and 8 in the EYT. Equation 11 was used to estimate SNP effects with year included as a fixed effect. The model was fit using the “solveRRBLUP” function from the AlphaMME R-package.

Estimated genetic values (EGV) were calculated using an additive model:

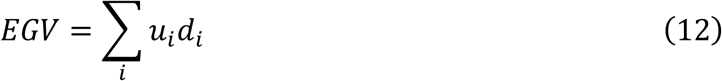

Where *u*_*i*_ is the SNP effect for locus, *i*, estimated via ridge regression (above), and *d*_*i*_ is the allele dosage value for an individual representing the expected number of alleles that they carry at a particular locus. This number will be an integer (e.g., 0, 1, or 2) for individuals genotyped on the SNP array, but may be a fractional value for individuals with inferred genotype dosages.

Lines were selected by choosing the lines with the highest genetic value.

### Headrow selection strategies

We considered five headrow selection strategies. Lines in the headrow were selected either at random (*random selection*), based on their observed phenotype by assuming a heritability of 0.05 for low-accuracy visual selection (*phenotypic selection*), using genomic selection with either their true genotype (*genomic selection*) or their inferred genotype dosages (*HTP-enabled genomic selection*), or using the high-dimensional phenotypes as a predictor for yield, (*phenomic selection* [11]). *Random selection* and *genomic selection* were included as lower and upper bounds on the accuracy of selection.

For *phenomic selection* we trained a model to predict a line’s estimated genetic value based on their high-throughput phenotypes. The training population used were the lines in the PYT. The model was fit using ridge regression (Equation 11), with the estimated genetic value as the dependent variable (from Equation 12), and the high-throughput phenotypes as the random effect design matrix. This model produced estimated effects for each high-throughput phenotype, which were used to estimate genetic values for non-genotyped lines. We also examined an alternative model where yield was used as the dependent variable, but found that this led to lower accuracies in all cases.

In the greenhouse-based headrow scenario, we did not consider *phenotypic selection*, under assumption that performance in the greenhouse would have low correlation with performance in the field, due to small plot sizes and large environmental differences between the greenhouse and production environments.

### Scenarios

We considered a total of 36 (2×2×3×3) scenarios based on the breeding program design (field-based or greenhouse-based headrows), species and number of chromosomes simulated (either 7 to represent barley or 21 to represent wheat), the heritability of all of the high-throughput phenotypes (between 0.1, 0.25, and 0.5), and the number of high-throughput phenotypes (either 25, 50, or 100). In each scenario we evaluated all five headrow selection strategies, and assumed a genetic map length of 100cM per chromosome, that the SNP array had 500 markers per chromosome, and all 500 lines in the PYT were used to estimate SNP effects.

We refer to the scenario with 100 high-throughput phenotypes, each with a heritability of 0.5 as the “best case-scenario”. This scenario is likely unrealistic given current high-throughput phenotyping technologies but represents the future potential of this technology.

In order to enable direct comparison between scenarios, we used a unified-burn across scenarios. In the unified burn-in we generated founder haplotypes for wheat and for barley and then simulated 20 years of phenotypic selection. The simulation was then saved at year 20,and this saved data was re-used as the starting point for each scenario. Using a unified burn-in reduced variation between the scenarios due to the position of the QTL effects, the simulation of founder haplotypes, and the initial rounds of pre-genomic selection.

We ran 10 replicates. In each replicate we re-simulated the unified burn-in and re-ran each scenario. Results were averaged across all replicates.

### Assessment

We assessed performance of the breeding program by tracking the mean genetic value of lines in the PYT, the prediction accuracy of the selection method in the headrow, and genotype accuracy for the inferred genotypes in the headrow.

Prediction accuracy was calculated as the correlation within a cross between the individuals’ estimated genetic values and their true genetic values. Predication accuracy was then averaged across crosses.

Genotype accuracy was measured as the correlation between an individual’s true genotype and their predicted genotype, corrected for their parent average genotype, and was averaged across individuals.

We used paired t-tests (paired within replicates) to evaluate whether the difference observed between scenarios and headrow selection strategies was significant.

## Results

In the parameter testing simulations we found that genotype accuracy was higher if more high-throughput phenotypes were used, if those phenotypes had higher heritability, and if the training population was larger. We also found that genotype accuracy decreased with an increasing size of the species genome. The results for these simulation are presented in Figure 1. In the breeding program simulations, we found that the inferred genotypes could be used to enable genomic selection on non-genotyped individuals and increase genetic gain compared to random selection, or in some scenarios phenotypic selection.

**Figure 1.**
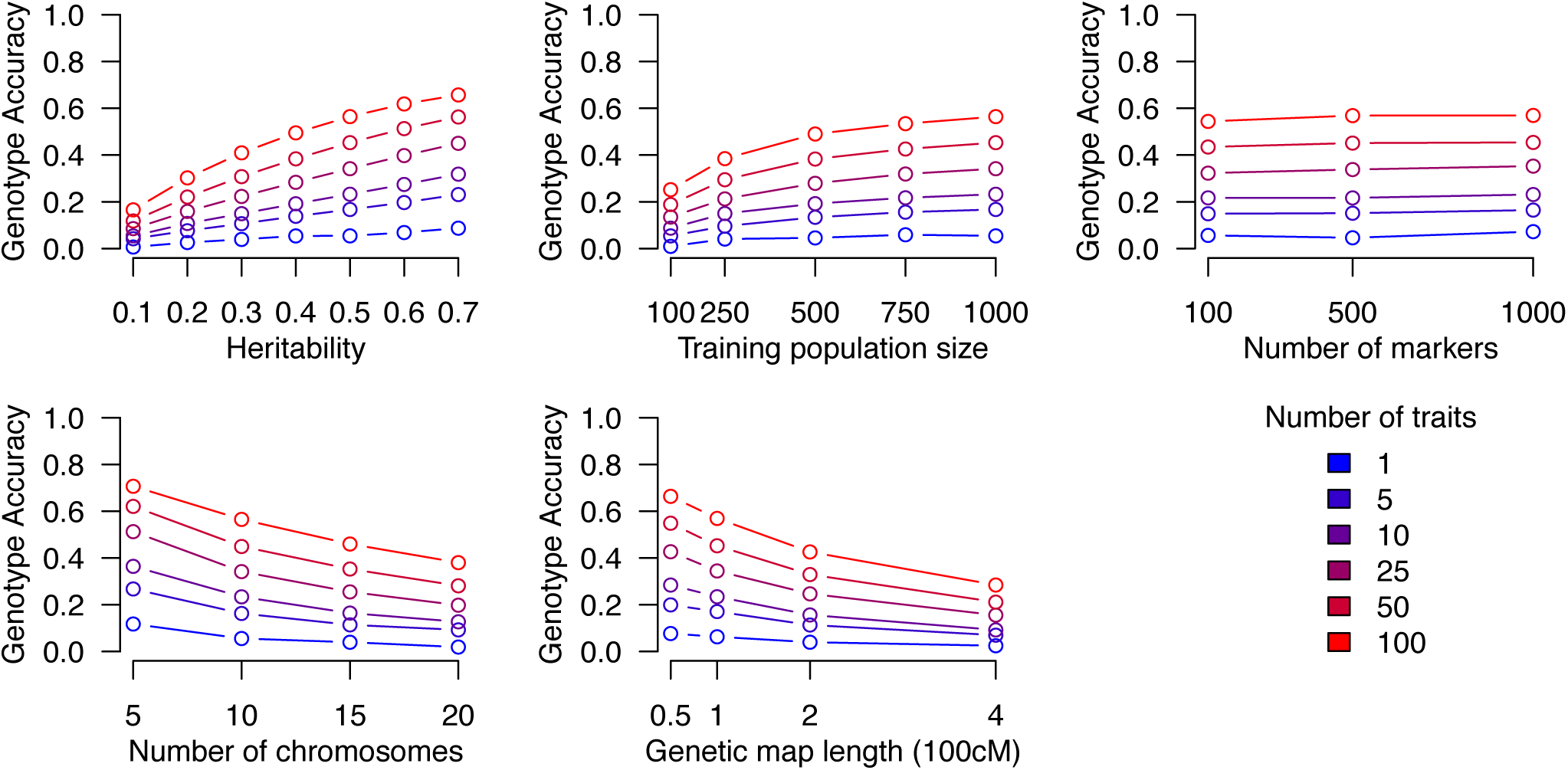
The influence of phenotype heritability, training population size, number of markers, number of chromosomes, and genetic map length on genotype accuracy.

### Parameter sensitivity analysis simulations

Genotype accuracy increased with larger numbers of high-throughput phenotypes measured, and higher heritability of the high-throughput phenotypes. For the number of high-throughput phenotypes measured, accuracy increased from 0.054 when 1 phenotype with a heritability of 0.5 was measured to 0.564 when 100 phenotypes with a heritability of 0.5 were measured. For the heritability of the phenotypes, accuracy increased from 0.166 when 100 phenotypes were used with a heritability of 0.1, to 0.564 when 100 phenotypes were used with a heritability of 0.5.

Increasing the training population size increased genotype accuracy. Accuracy was 0.252 when 100 individuals were used to estimate QTL effects. Accuracy increased to 0.490 when 500 individuals were used to estimate QTL effects, and increased again to 0.564 when 1,000 individuals were used to estimate QTL effects.

Increasing the number of chromosomes decreased genotype accuracy. Accuracy was 0.707 when the individual had 5, 100cM chromosomes. Accuracy decreased to 0.38 when the individual had 20, 100cM chromosomes. Increasing genetic map length also decreased genotype accuracy. Accuracy was 0.569 when the individual had 10, 100cM chromosomes. Accuracy decreased to 0.285 when the individual had 10, 400cM chromosomes.

### Breeding program simulations

In the breeding program simulations, we found that using *HTP-enabled genomic selection* in the headrow resulted in higher-genetic gain than *random selection*. Genetic gain was highest when the heritability of the high-throughput phenotypes was large, when more high-throughput phenotypes were used, and when barley (7 chromosomes) was considered compared to wheat (21 chromosomes).

### Best case scenario

In the best-case scenario we found that the genetic gain of *HTP-enabled genomic selection* was higher than *random selection* and *phenomic selection* in most cases for both wheat and barley, and higher than *phenotypic selection* in barley. In the best case scenario, we assumed that 100 high-throughput phenotypes were used and that each had a heritability of 0.5. We present the results for each selection method in Figure 2. Across all scenarios *random selection* led to the lowest genetic gain, and *genomic selection* led to the highest genetic gain.

**Figure 2.**
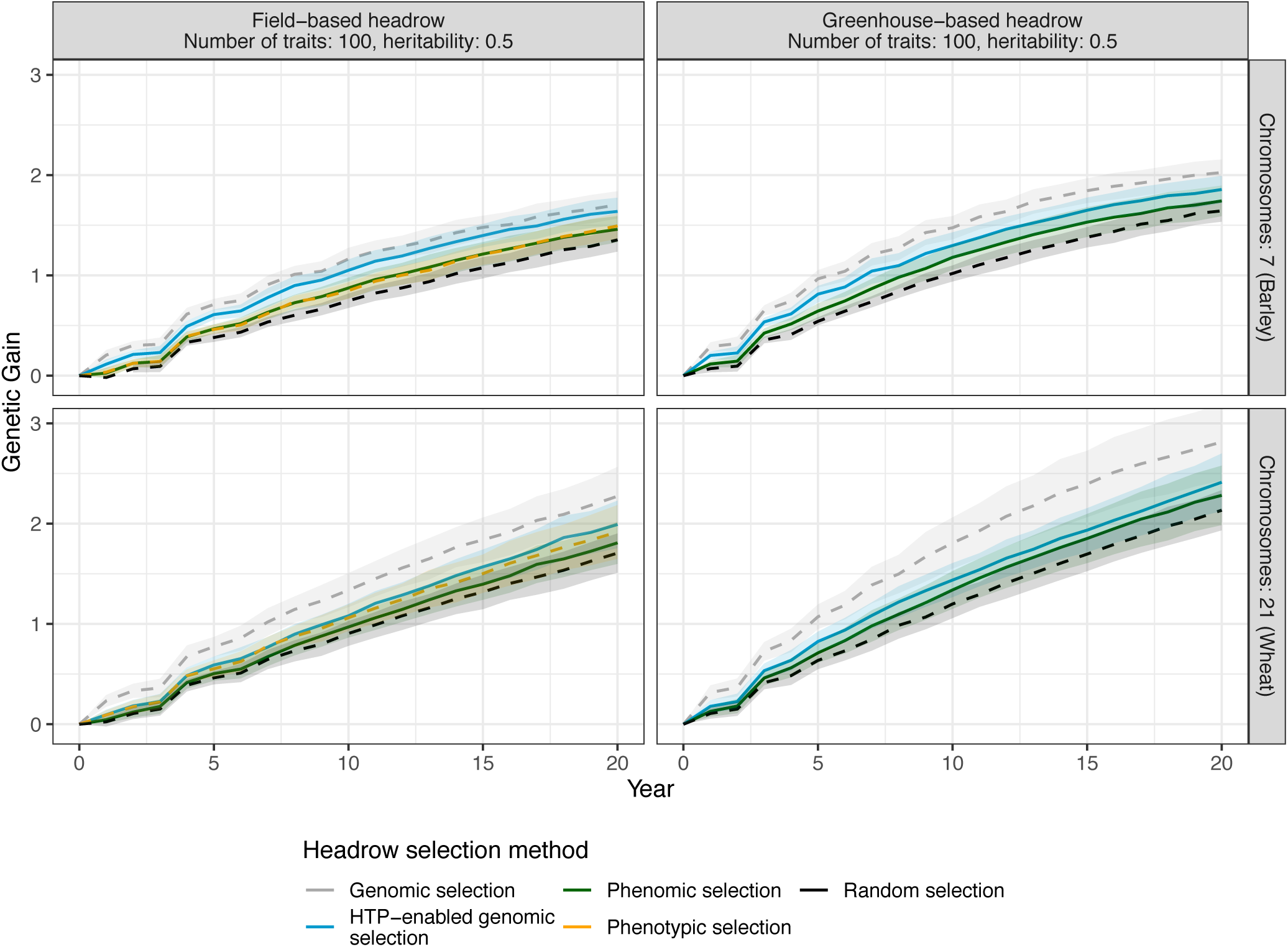
Genetic gain in the best-case scenario, represented as the average genetic value in the PYT at each year compared to year 0. All scenarios within the same species (barley or wheat) had the same 20 years of burn-in. For phenomic selection and HTP-enabled genomic selection, there were 100 high-throughput phenotypes, which each had a heritability of 0.5. Dashed lines represent indicate the selection strategies which did not use high-throughput phenotypes.

Using *HTP-enabled genomic selection* to select individuals in the headrow, led to a larger increase in genetic gain compared to *phenomic selection* in most scenarios. In barley, the genetic gain using *HTP-enabled genomic selection* was significantly higher than *phenomic selection* in the field-based headrow scenario (paired t-test, t(9) = 5.58, p<0.01), but was only marginally significantly higher in the greenhouse-based headrow scenario (paired t-test, t(9) = 2.12, p=0.06). In wheat, the genetic gain using *HTP-enabled genomic selection* was significantly higher than *phenomic selection* in both the field-based headrow scenario (paired t-test, t(9) = 3.85, p<0.01), and the greenhouse-based headrow scenario (paired t-test, t(9) = 2.76, p=0.02).

Using *HTP-enabled genomic selection* led to a higher genetic gain then *phenotypic selection* in the field-based headrow scenario in barley (paired t-test, t(9) = 7.55, p<0.01), but not in wheat (paired t-test, t(9) = 2, p=0.08). We did not evaluate the performance of *phenotypic selection* in greenhouse-based headrow scenario, due to the fact that the greenhouse environment is likely sufficiently different from the field environment, making phenotypic selection unreliable in practice.

### Genetic gain based on number of high-throughput phenotypes and their heritabilities

In all scenarios, the genetic gain obtained using *HTP-enabled genomic selection* was higher than that obtained using *random selection*, but lower than that of *genomic selection*. The relative performance of *HTP-enabled genomic selection* compared to *phenotypic selection* depended on the scenario. The results of these scenarios are presented in Figure 3

**Figure 3.**
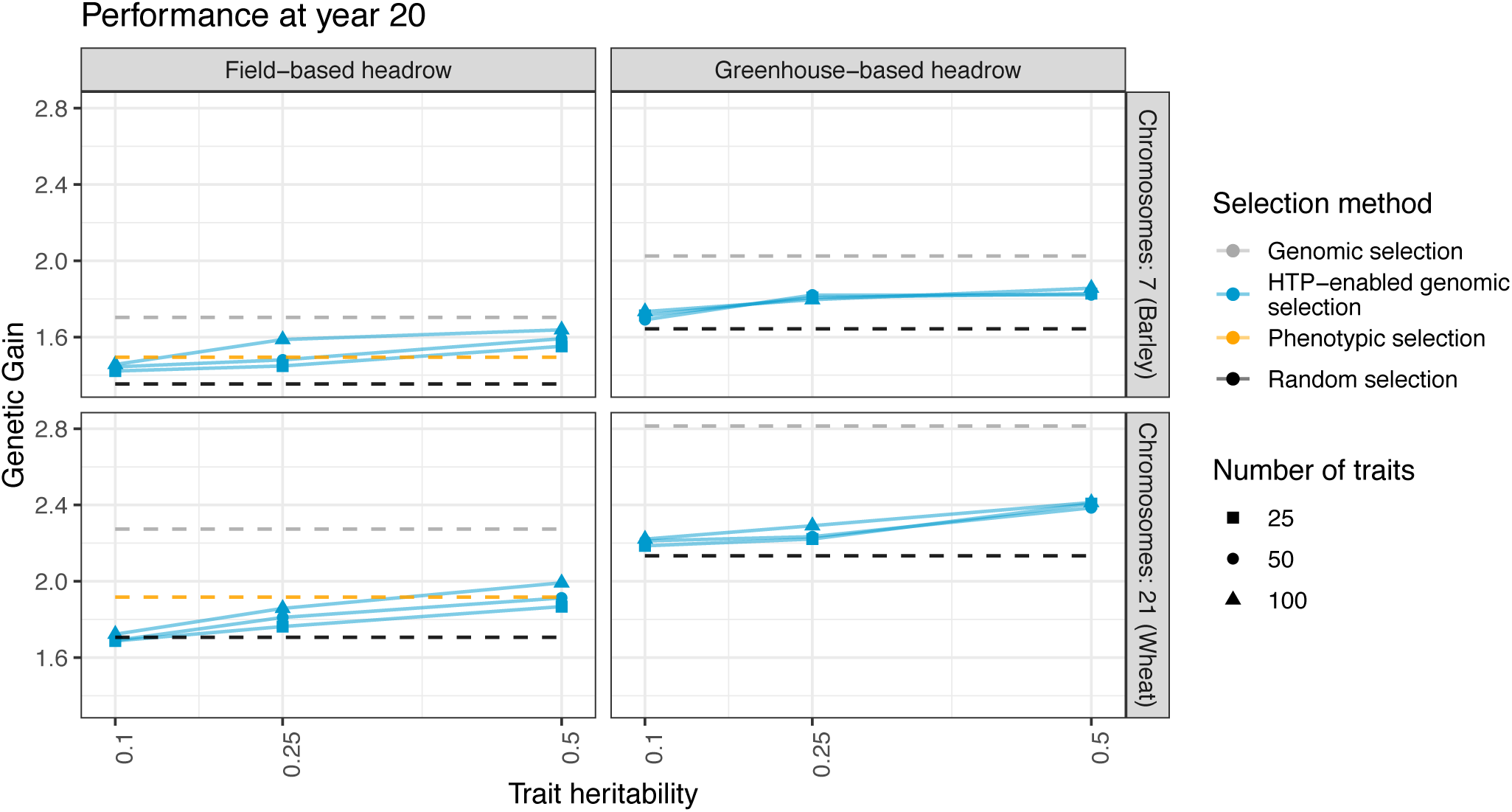
Genetic gain compared to year 0 in the final year of the breeding program. All scenarios within the same species (barley or wheat) had the same 20 years of burn-in.

In barley, *HTP-enabled genomic selection* led to higher genetic gain compared to *phenotypic selection* when 50 or 100 high-throughput phenotypes were used with a heritability of 0.5, or when 100 high-throughput phenotypes were used with a heritability of 0.25. In wheat, *HTP-enabled genomic selection* never significantly outperformed *phenotypic selection*.

In addition, we found that the greenhouse-based headrow scenario had higher genetic gain than the field-based headrow scenario when both random selection (barley: paired t-test, t(9) = - 10.02, p<0.01, wheat: paired t-test, t(9) = −14.62, p<0.01) and genomic selection were used (barley: paired t-test, t(9) = −6.36, p<0.01, wheat: paired t-test, t(9) = −7.84, p<0.01).

### Relationship between genotype accuracy and prediction accuracy

We found a linear relationship (r^2^=0.696) between genotype accuracy, and the ratio of prediction accuracy when the true genotypes were used to the prediction accuracy when the inferred genotypes were used. The results of this comparison are presented in Figure 4.

**Figure 4.**
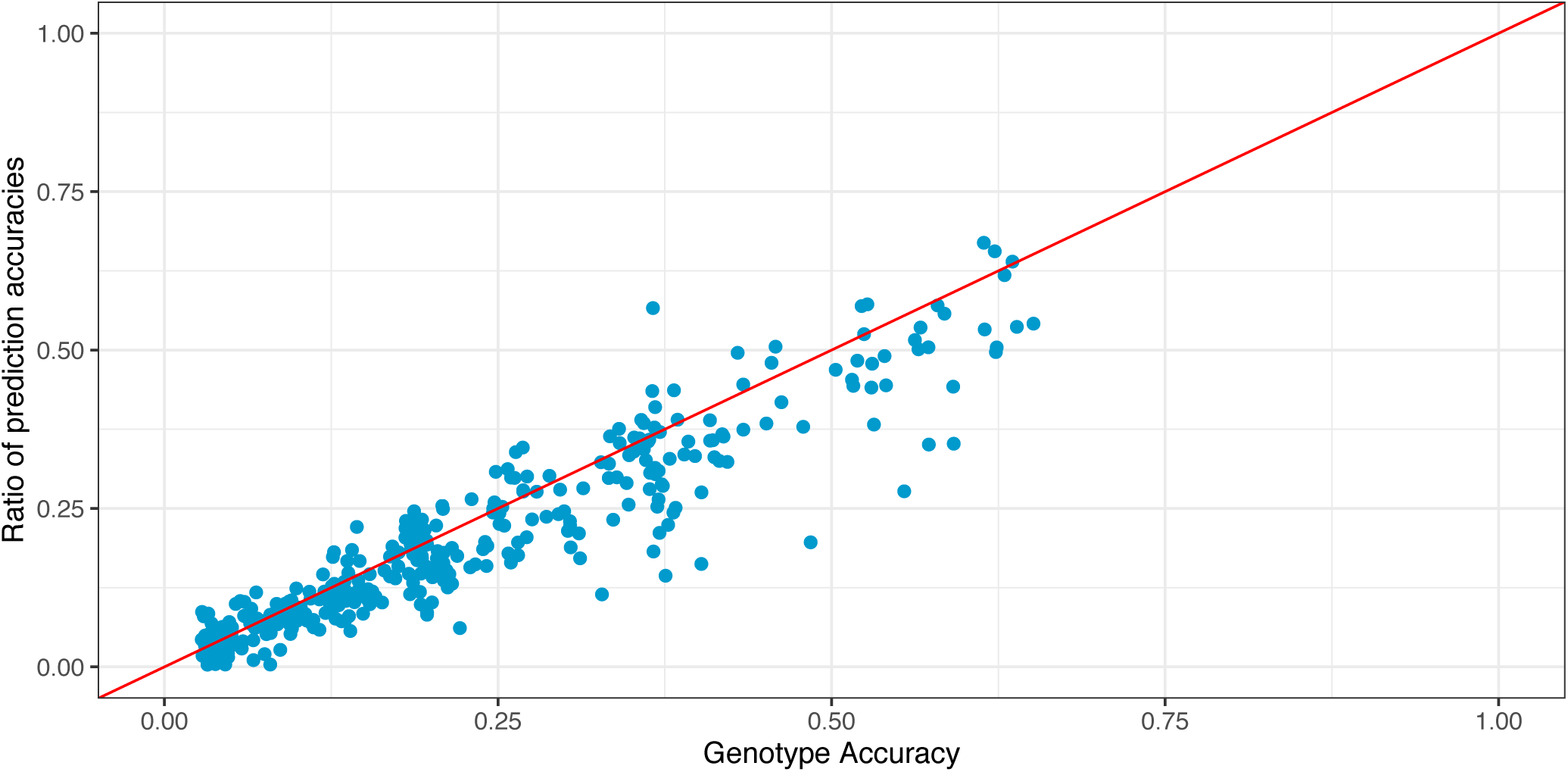
Comparison between genotype accuracy and prediction accuracy. Genotype accuracy is measured as the correlation between an individual’s true and inferred genotypes. The ratio of prediction accuracies is calculated as prediction accuracy using HTP-enabled genomic selection divided by prediction accuracy using genomic selection. Results are combined across all scenarios and replicates. The red line is the y=x line.

## Discussion

In this paper we present a novel approach for using high-throughput phenotype data to infer the genotypes of an individual. We conducted two sets of simulations. In the first set of simulations we found that accuracy was higher if more high-throughput phenotypes were used and if those phenotypes had higher heritability. We also found that genotype accuracy decreased with an increasing size of the species genome. In the second set of simulations, we analysed whether this method could be used to increase genetic gain in a plant breeding program, by enabling genomic selection on non-genotyped individuals in the headrow. We found that using high-throughput phenotype enabled genomic selection, we could obtain higher genetic gain than random selection, and in some cases phenotypic selection. In the remainder of the discussion, we discuss the factors that influenced the accuracy of the inferred genotypes, highlight the differences between this approach of using high-throughput to increase the accuracy of selection to alternative approaches, and discuss where this method might be most advantageous in modern breeding programs.

### Factors that influence genotype accuracy

In our simulation study, we found that accuracy depended on both the high-throughput phenotypes which can be changed by selecting alternative sets of phenotypes, and the genome of a species, which cannot be changed.

With regards to the high-throughput phenotypes measured, our main finding is that genotype accuracy increased substantially with more of high-throughput phenotypes, and with higher heritability of those phenotypes. We found that moderate genotype accuracy (∼0.5) could be reached with 100 high-throughput phenotypes, each with a heritability of 0.5. This scenario formed the basis of our “best case” scenario in the breeding program simulations. It was found to achieve a genetic gain in barley higher than that of phenotypic selection, and in wheat, equivalent to that of phenotypic selection. Based on current technology it may be possible to collect high-throughput phenotypes of this density in the near future. Existing high-throughput phenotyping methods are able to produce a moderate number of high-dimensional phenotypes which show moderate heritability [8,11]. The number of phenotypes collected can be increased by collecting the phenotypes at multiple time points over the course of the growing cycle [3,10], although there may be diminishing returns if the resulting phenotypes are correlated, or by collecting phenotypes on multiple parts of the plants, e.g., both on the leaves and the grains [11].

We found that the number of markers on the high-density SNP array had no impact on genotype accuracy, and that training population size had only a limited impact on genotype accuracy. These findings likely result from the fact that the method attempts to infer entire inherited haplotype blocks instead of trying to infer an individual genotype. For SNP array density, accuracy depended primarily on whether the model was able to detect which haplotypes the individual inherited from their parents. If the shared haplotype segments were detected correctly, accuracy will be high no matter the density of the SNP array. For training population size, estimating the genetic values of an entire haplotype block is easier than estimating the genetic values for any particular marker, allowing for smaller training populations to be used. There was also an asymptotic plateau of accuracy depending on training population size, suggesting that although small training population sizes may yield low accuracy genotypes, there is a limit to how much accuracy can be increased by simply increasing the training population size. In addition, even though small training populations can be used, building training population size may be an issue if models need to be re-trained each year/location to account for environmental and temporal differences in the phenotype expression [2].

In regards to the genome of the species, we found that genotype accuracy was lower in species with a longer genetic map length, and a larger number of chromosomes. Both of these factors increase the number of recombinations that might occur between an individual and their parents, making the inference problem more challenging. Unlike the number of traits which can be increased by technological advances, the genetic architecture of a species is fixed. We find that in the context of our genetic gain simulations, the highest gains were found when simulating a barley breeding program (7 chromosomes), compared to a wheat breeding program (21 chromosomes).

In addition to the species, the method of generating inbred individuals may also impact the effective genetic map length of the species. Applying this algorithm to individuals generated via multiple rounds of selfing will likely produce lower genotype accuracies due to the larger number of meiosis separating an individual from its parents compared to an individual produced via doubled haploid technologies. In addition, it may be possible to apply this approach to outbred individuals if the parent’s genotypes are known and phased (particularly of importance for livestock and some crop species), but accuracy will be lower due there being twice as many effective haplotype segments.

### Using high-throughput phenotypes for genomic selection compared to previous approaches

Using high-throughput phenotype data to infer genotypes and enable genomic selection offers a radically different way to use high-throughput phenotype data. Past approaches for utilizing high-throughput phenotypes to increase the accuracy of selection have primarily focused on their ability to provide a proxy for a phenotype of interest [2,7], increase prediction accuracy as correlated traits [3], or estimate the genetic covariance between individuals, i.e., phenomic selection [11]. We compare the method presented here with each of these alternatives in turn.

Compared to past work that has used high-throughput phenotypes as proxies for selection targets, our method of using high-throughput phenotypes to enable genomic selection offers the potential for higher cross-environment predictions, and ability to select on selection targets that are uncorrelated with the phenotypes collected. Past work has found that many high-throughput phenotypes e.g., NVDI, can be used as a proxy for yield or drought tolerance [3,7]. However, the presence of genotype-by-environment interactions may limit the predictive ability of these models to the environments in which the high-throughput phenotypes are collected. This means that using high-throughput phenotypes as a proxy for the selection target should be used when the plants are grown in an environment similar to the production environment, and when there is a strong correlation between the high-throughput phenotypes and the selection target. In comparison, the method presented here produces inferred genotypes, which can be used to provide genomic predictions independent of both the growing environment and the selection target. This may be particularly useful if the growing environment of an individual is dissimilar to the production environment, or if the selection target has low correlation with the high-throughput phenotypes measured.

Another way to use high-throughput phenotype data is to use individual phenotypes as correlated traits in a genomic prediction model for genotyped individuals [10]. The method presented here offers a complimentary way to use high-throughput phenotypes on non-genotyped individuals. There is a synergy between these methods, since the high-throughput phenotype data for the training population can be used twice: first to increase the accuracy of selection in the training population itself, and second to allow for genomic selection to be applied to non-genotyped individuals grown in the same field.

Phenomic selection [11] offers the most similar approach to the method presented here. In phenomic selection where a large number of high-throughput phenotypes are used as a stand-in for genetic markers to estimate the relatedness between individuals. In our simulations, we evaluated both methods, and directly inferring the individual’s genotypes led to higher prediction accuracy than performing phenomic selection. This result likely is a result of the fact none of the high-throughput phenotypes correlated with the selection targets in these simulations, and phenomic selection is able to use correlations with selection targets to increase accuracy. In practice, many of the high-throughput phenotypes will correlate with the selection target (e.g., NVDI correlates with yield) which may increase the accuracy of phenomic selection. Including correlated traits into the genomic prediction models used for high-throughput phenotype enabled genomic selection is an area of future research.

### Applying high-throughput phenotype enabled genomic selection in a breeding program

The primary motivation of this work was to develop a method that allowed genomic selection to be performed on non-genotyped individuals. In our simulations, we found that using high-throughput phenotypes to enable genomic selection could increase genetic gain compared to random selection, and in some cases phenotypic selection.

When applying high-throughput phenotype enabled genomic selection to breeding programs, the lower accuracies produced by this method will be least useful in situations where the breeder is able to easily select non-genotyped individuals via visual selection, or another proxy. In all wheat scenarios, we found that phenotypic selection either outperformed, or performed similarly to high-throughput enabled genomic selection. In barley, phenotypic selection underperformed selecting individuals with inferred genotypes, particularly when the number of phenotypes is large. Situations where this is likely to be the case, are cases when individuals are grown in an environment similar to the target environment, and either visual selection is accurate, or there are high-accuracy proxies for phenotypic selection (e.g., using high-throughput phenotypes such as NVDI).

Using high-throughput phenotypes to enable genomic selection will be most useful in cases where the breeder is unable to accurately select non-genotyped individuals. These will be cases where phenotypic selection has either low accuracy, or if the selection targets cannot be easily measured, e.g., grain yield.

One particular place where this method would be advantageous is to enable genomic selection to occur in a greenhouse environment, and thereby reduce the cycle time of the breeding program. Traditionally, it is hard to make selection decisions in the greenhouse environment given the small plot sizes and environmental differences between the growing environment and the production environment. Genomic selection can overcome this issue by allowing selections decisions without directly assessing the individual’s phenotype. Our method enables the use of genomic selection (albeit at lower accuracy) by collecting high-throughput phenotypes instead of genotyping them. This would allow a higher genetic gain, by coupling reduced cycle time, with a cost-effective selection strategy. In addition, the greenhouse environment may also be ideal for automating the collection of high-throughput phenotypes on individuals, particularly for phenotypes measured at multiple time points or on multiple tissues.

## Conclusions

In this paper we develop and test a method which uses high-throughput phenotypes to infer the genotypes of an individual. The inferred genotypes can then be used to perform genomic selection. This method presents a radically different way to use high-throughput phenotype data to enable genomic selection. In simulations, we found that it was possible to use these genotypes to increase the genetic gain in a plant breeding program. Although this method requires a large number of heritable phenotypes to obtain high-accuracy genotypes, we believe these phenotypes will become increasingly available as phenotyping technologies improve, allowing this method to enable genomic selection on massive numbers of individuals at a low cost.

## Author contributions

AW designed the genotype inference algorithm. AW, CG, JH designed the simulations. AW ran the simulations and analyzed the results. All authors contributed to writing the manuscript and approved the final manuscript

## Acknowledgements

This work has made use of the resources provided by the Edinburgh Compute and Data Facility (ECDF) (http://www.ecdf.ed.ac.uk).

